# Exploiting NMR Ensemble Heterogeneity Enables Small Molecule Discovery Against Dynamic Protein-Protein Interfaces

**DOI:** 10.64898/2026.02.04.703700

**Authors:** Hossam Nada, Sungwoo Cho, Ashraf N. Abdo, Moustafa Gabr

## Abstract

Protein-protein interactions governed by conformationally heterogeneous domains remain difficult to drug because ligand-competent states are often absent from single static structures. Here, we present AtlasNMR, a statistical framework that transforms multi-model NMR ensembles into screening-ready conformational hypotheses for small molecule discovery. Using the neuronal nitric oxide synthase (nNOS) PDZ domain that engages the adaptor protein CAPON (NOS1AP) as a model system, AtlasNMR identified two representative conformational states capturing the dominant and minor populations of the NMR ensemble. Ensemble-based virtual screening followed by consensus ranking yielded **MC-3**, a small molecule modulator that disrupts the NOS1-NOS1AP interaction in live cells and directly engages the nNOS PDZ domain. **MC-3** produced convergent neuroprotective effects in disease-relevant neuronal models by reducing amyloid-β-induced cytotoxicity, suppressing NMDA-driven nitrosative stress, and attenuating pathological tau phosphorylation, while exhibiting a balanced early lead-like ADME and safety profile. Together, this work establishes a generalizable strategy for exploiting NMR ensemble heterogeneity to enable small molecule discovery against dynamic protein-protein interfaces.

## 1. Introduction

Protein-protein interactions (PPIs) mediated by conformationally dynamic domains remain among the most challenging targets for small molecule discovery. Ligand-competent states are often transient, weakly populated, or inaccessible in single static structures, limiting the effectiveness of conventional structure-based screening approaches. For intracellular signaling hubs that rely on modular interaction domains, this conformational plasticity represents a fundamental barrier to chemical intervention and calls for strategies that explicitly account for ensemble heterogeneity.

The carboxy-terminal PDZ ligand of neuronal nitric oxide synthase (CAPON) is a key regulatory protein that modulates nitric oxide (NO) signaling in various physiological and pathological processes^1,2^. CAPON has emerged as a compelling therapeutic target due to its involvement in multiple disease pathways, particularly in Alzheimer’s disease and other neurodegenerative disorders where its modulation could provide critical neuroprotection^3-5^. However, targeting the CAPON/nNOS interaction presents significant challenges that have thus far prevented successful drug development.

Nitric oxide synthase (NOS) exists in three distinct isoforms: neuronal-type (nNOS), inducible-type (iNOS), and endothelial-type (eNOS)^6^. Each isoform serves specialized functions across different tissues including the cerebellum, skeletal muscles, kidneys, blood vessels, and immune cells^7-9^. Among the nine known proteins that interact with nNOS, CAPON acts as a crucial regulator of nNOS activity through its unique ability to compete with postsynaptic density protein 95 (PSD-95) for binding to the nNOS PDZ domain^10^. This competitive interaction fundamentally alters NO production and downstream signaling cascades, positioning CAPON as a master regulator of nitric oxide homeostasis.

The CAPON signaling pathway operates through a complex molecular switching mechanism that directly impacts neuronal function and cellular metabolism. Under normal conditions, nNOS forms a functional complex with N-methyl-D-aspartate (NMDA) receptors and PSD-95, facilitating calcium-dependent NO production in response to glutamatergic signaling^11^. However, CAPON disrupts this canonical NMDAR-PSD95-nNOS complex by competing for the same PDZ-binding motif on nNOS, resulting in the formation of an alternative NMDAR-CAPON-nNOS complex^2,12^. This molecular rearrangement significantly attenuates nNOS activity and reduces NO production.

The conformational heterogeneity observed in nNOS is biologically significant and likely plays a crucial role in CAPON binding. CAPON, which contains a C-terminal PDZ-binding motif, likely engages with multiple nNOS conformational states to achieve its regulatory function. The presence of distinct conformational clusters suggests that nNOS exists in a dynamic equilibrium between multiple states, consistent with the conformational selection model wherein CAPON selectively binds and stabilizes specific conformations that favor interaction^13^. This plasticity may be essential for facilitating the formation of the nNOS-CAPON complex, which plays a critical role in neuronal signaling by anchoring nNOS to specific subcellular locations^14^.

Inhibiting the CAPON/nNOS interaction presents a critical therapeutic opportunity for neuroprotection in Alzheimer’s disease and other neurodegenerative conditions^15,16^. By preventing CAPON from binding to nNOS, the pathological redistribution of nNOS can be avoided, reducing excessive NO production and the subsequent excitotoxicity and oxidative stress, two key pathological mechanisms in neurodegeneration^17-19^. Disruption of the CAPON/nNOS complex has the potential to restore proper nNOS regulation through its interaction with PSD-95^20^, preventing the deleterious effects of aberrant NO signaling and protecting neurons from excitotoxic damage^21,22^. This represents a promising neuroprotective strategy that could slow or prevent neuronal loss in Alzheimer’s disease.

Herein, we report a strategy for systematically leveraging NMR ensemble heterogeneity to enable small molecule discovery against dynamic protein-protein interfaces and demonstrate its application through the identification and biological validation of **MC-3**, a neuroprotective modulator of the CAPON/nNOS interaction.

## 2. Results and discussion

### 2.1. AtlasNMR: mapping NMR ensemble heterogeneity into screening-ready chemical hypotheses

Protein conformational heterogeneity presents a major barrier to structure-based discovery against protein-protein interaction interfaces, particularly when ligand-competent states are weakly populated or obscured within multi-model structural data. AtlasNMR (Figure 1) was developed to address this challenge by providing a statistically grounded framework that converts NMR ensemble heterogeneity into a minimal set of representative conformational states suitable for ensemble-based virtual screening and downstream chemical decision-making.

**Figure 1.**
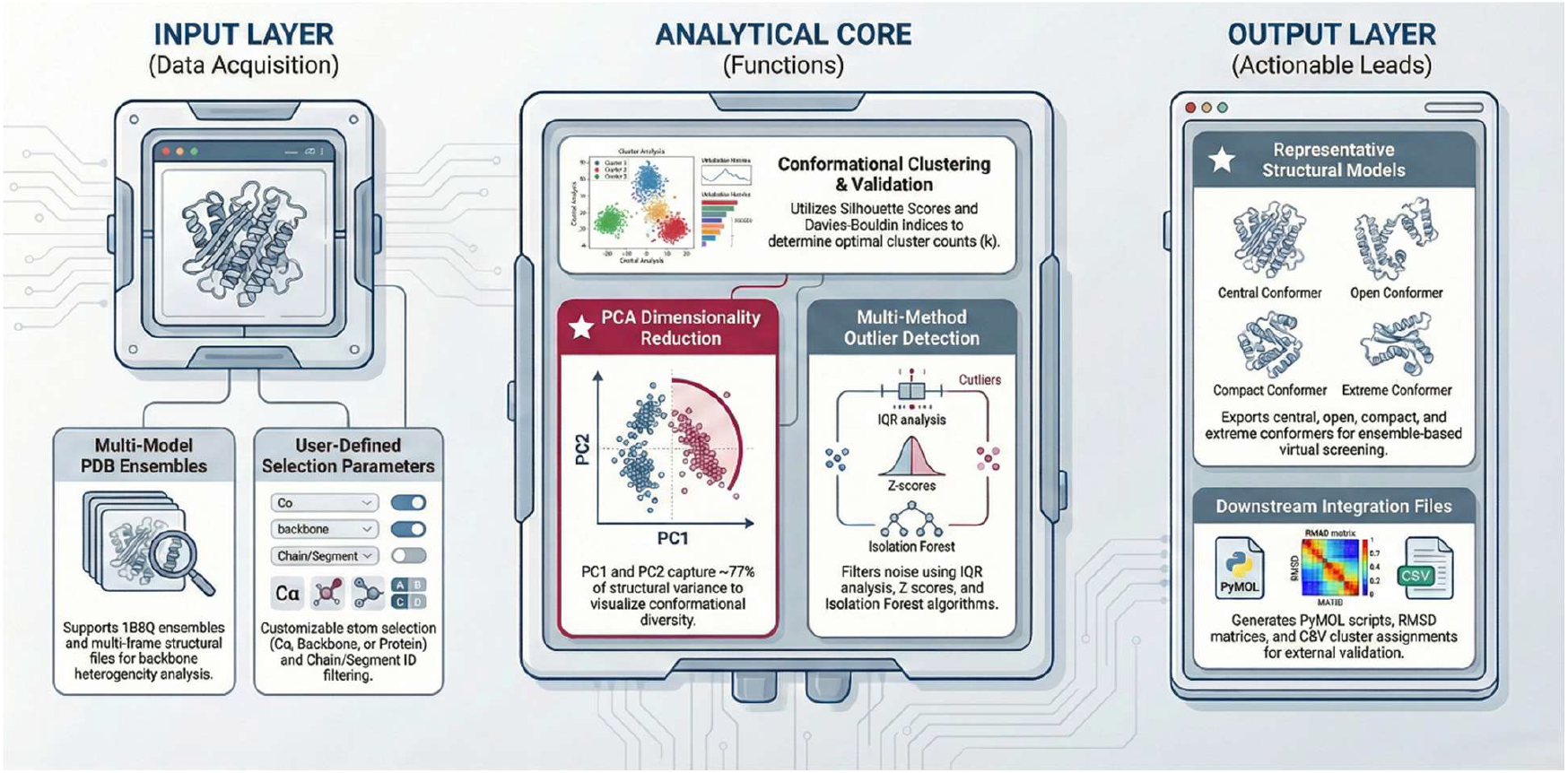
AtlasNMR framework for converting NMR ensemble heterogeneity into screening-ready conformational hypotheses. Multi-model NMR structures are aligned and analyzed using statistical outlier detection and clustering to identify dominant and minor conformational states. Representative structures capturing maximal ensemble diversity are selected and used for ensemble-based virtual screening, enabling small molecule discovery against dynamic protein–protein interaction interfaces.

AtlasNMR analyzes multi-model NMR structures through a sequential workflow that integrates structural alignment, statistical outlier detection, conformational clustering, and representative selection. All ensemble members are first superimposed using least-squares fitting of backbone atoms to remove rotational and translational variance prior to quantitative comparison. Pairwise root mean square deviation (RMSD) matrices are then computed to capture the full conformational landscape sampled by the ensemble. This quantitative representation enables objective assessment of structural similarity and diversity across all models.

To ensure robust identification of conformational states, AtlasNMR employs a consensus outlier detection strategy that integrates three independent approaches: interquartile range (IQR) analysis, the Isolation Forest algorithm, and hierarchical clustering with Ward’s linkage. Models identified as outliers by at least two methods are excluded from representative selection, preventing rare or artifactual conformations from biasing downstream screening efforts. This step is critical for maintaining chemical interpretability and avoiding overfitting to non-physiological states.

Following outlier filtering, AtlasNMR systematically evaluates candidate cluster numbers using multiple orthogonal quality metrics, including silhouette coefficients, the Davies-Bouldin index, and the Calinski-Harabasz score. For the nNOS PDZ domain ensemble, these metrics converged on a two-cluster solution (k = 2), revealing a dominant conformational population and a minor but structurally distinct alternative state. This bimodal distribution provides a chemically meaningful representation of the ensemble, consistent with conformational selection mechanisms at dynamic protein-protein interaction interfaces.

Beyond clustering, AtlasNMR identifies representative structures using a consensus voting strategy that integrates medoid selection, centroid proximity, minimax optimization, and density-based criteria. This multi-criteria approach ensures that selected representatives capture both the central tendency and maximal structural diversity of each cluster while maintaining comparable surface topology and solvation characteristics. Principal component analysis (PCA) further confirms that the selected representatives occupy extreme and well-separated regions of conformational space, validating their suitability for ensemble-based docking.

Importantly, AtlasNMR is designed to support chemical decision-making rather than exhaustive structural description. By reducing complex NMR ensembles into a small number of statistically justified, screening-ready conformations, the framework enables practical incorporation of experimentally observed protein dynamics into virtual screening workflows. This approach provides a generalizable path for exploiting NMR-derived conformational heterogeneity in small molecule discovery against dynamic protein-protein interaction interfaces.

### 2.2. NMR analysis of NOS/CAPON interaction domain

AtlasNMR was employed to analyze the NMR structure of NOS/CAPON (PDB ID: 1B8Q). Cross-validation analysis employing the silhouette coefficient method conclusively demonstrates that the nNOS PDZ domain NMR ensemble optimally partitions into two distinct conformational clusters (k=2, silhouette score = 0.52). This bimodal distribution suggests the presence of two functionally relevant conformational states in solution. The silhouette score of 0.52 (Figure S1A) indicates good cluster separation and cohesion, validating the biological relevance of this clustering scheme. Multiple independent clustering quality metrics corroborate this finding. The Davies-Bouldin Index, which measures the average similarity ratio of each cluster with its most similar cluster, reaches its optimal value at k=2, indicating maximal inter-cluster separation. Similarly, the Calinski-Harabasz Score, a ratio of between-cluster to within-cluster variance, exhibits its highest value at k=2, further supporting the two-state model. This convergence of multiple statistical measures provides robust evidence for the selection of a two-cluster representation.

Analysis of clusters reveal a highly asymmetric population distribution (Figure 2 and S1), with Cluster 0 (C0) comprising approximately 87% of conformers (n=13) and Cluster 1 (C1) representing 13% (n=2). This distribution pattern is consistent with a dominant conformational state and a minor, yet structurally distinct, alternative conformation. In the context of drug discovery, this population bias suggests that C0 represents the thermodynamically favored state likely encountered by potential ligands under physiological conditions, while C1 may represent a transient or induced-fit conformation relevant to protein-peptide recognition.

**Figure 2.**
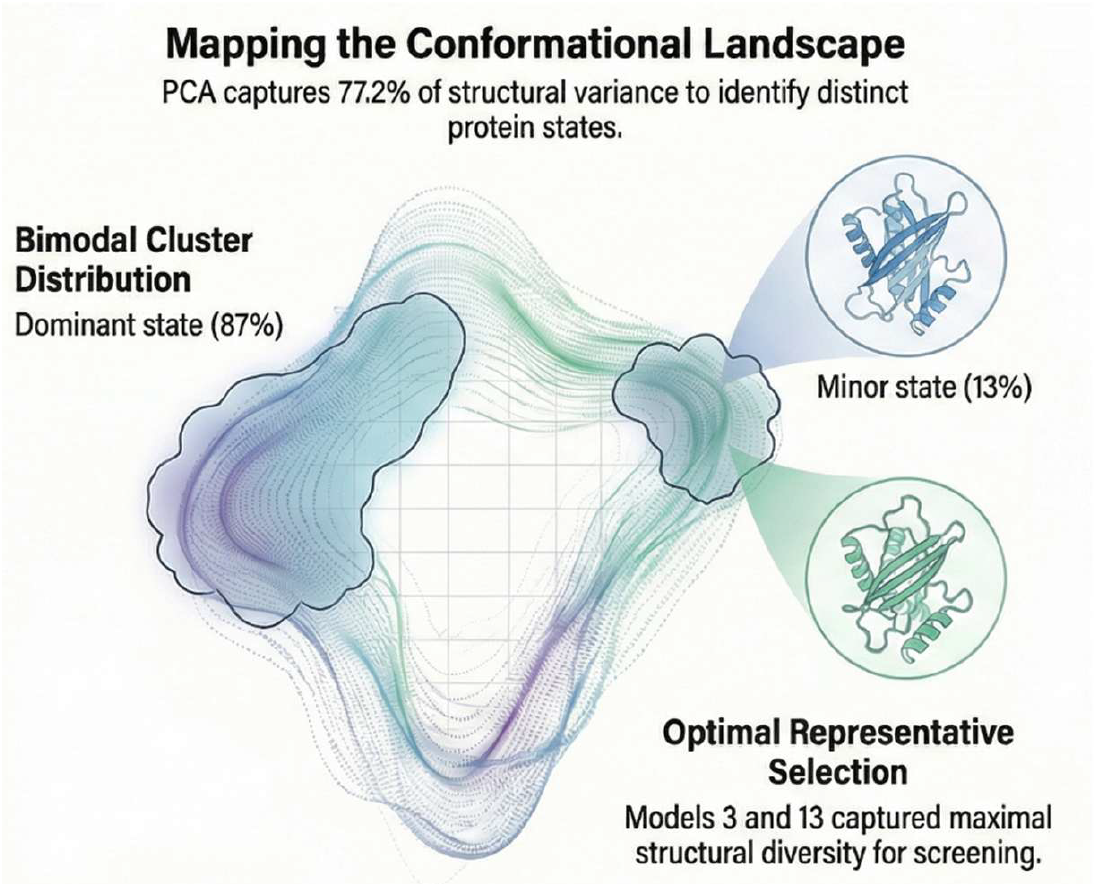
Conformational landscape of the nNOS PDZ domain derived from NMR ensemble analysis. Principal component analysis (PCA) of the aligned NMR ensemble (PDB ID: 1B8Q) reveals a bimodal conformational distribution corresponding to two structurally distinct states. Cluster 0 represents the dominant conformational population, while Cluster 1 corresponds to a minor but clearly separated alternative state. Representative structures selected from each cluster (models 3 and 13) occupy extreme and well-separated regions of conformational space, capturing the maximal structural diversity of the ensemble. This analysis establishes a chemically meaningful basis for ensemble-based virtual screening by identifying discrete conformational states that would be obscured in single-structure representations.

Principal component analysis (PCA) reveals that the first two principal components account for 77.2% of the total structural variance, indicating that the conformational heterogeneity can be effectively captured in a low-dimensional space. This high cumulative variance explained by the first two PCs validates the use of dimensionality reduction for ensemble analysis and confirms that the major conformational transitions occur along well-defined collective coordinates. The PCA-based cluster visualization demonstrates clear spatial separation between C0 and C1 along PC1, with representatives 3 and 13 positioned at opposite extremes of this primary axis of variation. This positioning indicates that these structures capture the maximal conformational diversity present in the ensemble. Representative 3 occupies the densely populated region of conformational space corresponding to C0, while representative 13 samples the less frequently visited but structurally distinct region corresponding to C1. The spatial distribution in the three-dimensional PCA space (PC1-PC3) further confirms the discrete nature of these conformational states, with C0 forming a compact cluster and C1 exhibiting greater conformational flexibility.

Lastly, analysis of surface-accessible residues (Figure S1C) across the ensemble reveals modest variation (55-62 residues), with a mean of approximately 59 residues. This relatively narrow distribution indicates consistent surface topology despite conformational heterogeneity, suggesting that the observed structural variance primarily involves side-chain reorientations and local backbone adjustments rather than large-scale domain movements. Representatives 3 and 13 exhibit surface residue counts within one standard deviation of the ensemble mean, confirming their structural representativeness. The conservation of surface area across conformers has important implications for virtual screening, as it ensures comparable solvation energetics and surface complementarity calculations. Structures with extreme surface area deviations could introduce systematic biases in scoring functions; the absence of such outliers in the selected representatives validates their suitability for docking studies.

### 2.3. Consensus Virtual screening

To identify small molecules capable of disrupting the CAPON-NOS interaction, an in-house Enamine library comprising 30,000 compounds was subjected to consensus virtual screening targeting the CAPON-NOS binding interface. The screening campaign utilized two representative protein conformations (models 3 and 13) to account for structural flexibility at the binding site. Virtual screening was performed using the GLIDE module of Schrödinger through a three-stage hierarchical docking protocol consisting of High-Throughput Virtual Screening (HTVS), Standard Precision (SP), and Extra Precision (XP) modes. A stepwise enrichment strategy was implemented in which only the top 10% of compounds from each stage progressed to the subsequent docking level, ensuring progressive refinement of hit quality.

To maximize prediction reliability, an ensemble consensus scoring pipeline integrating five complementary ranking algorithms was employed: z-score normalization, percentile ranking, weighted averaging (0.4 Glide/0.6 MMGBSA), Pareto frontier analysis, and dual-cutoff filtering. This multi-criteria approach identified 29 Pareto-optimal compounds occupying the non-dominated frontier, representing hits that simultaneously optimize both Glide docking scores and MMGBSA ΔG binding energies. These compounds were prioritized for experimental validation based on their superior predicted binding characteristics.

### 2.4. Identification of MC-3 as a potential NOS1-NOS1AP interaction inhibitor

Twenty-nine hit compounds were identified through consensus virtual screening and subsequently evaluated in a single-dose NanoBRET assay (Figure 3) to assess their ability to disrupt the NOS1-NOS1AP (CAPON) protein-protein interaction. NanoBRET, a bioluminescence resonance energy transfer technique, enables real-time monitoring of protein-protein interactions in live cells with high sensitivity and specificity. Based on their activity profiles and predicted binding properties to the NOS1 PDZ domain, three compounds (Z968614410, Z2607801368, and Z1204854808) were selected for further characterization. Dose-response analysis revealed that all three compounds induced concentration-dependent disruption of the NOS1-CAPON complex, confirming their functional activity in a cellular context. These findings validate the computational predictions and establish these compounds as promising lead candidates for further optimization and development as NOS1-CAPON interaction inhibitors.

**Figure 3.**
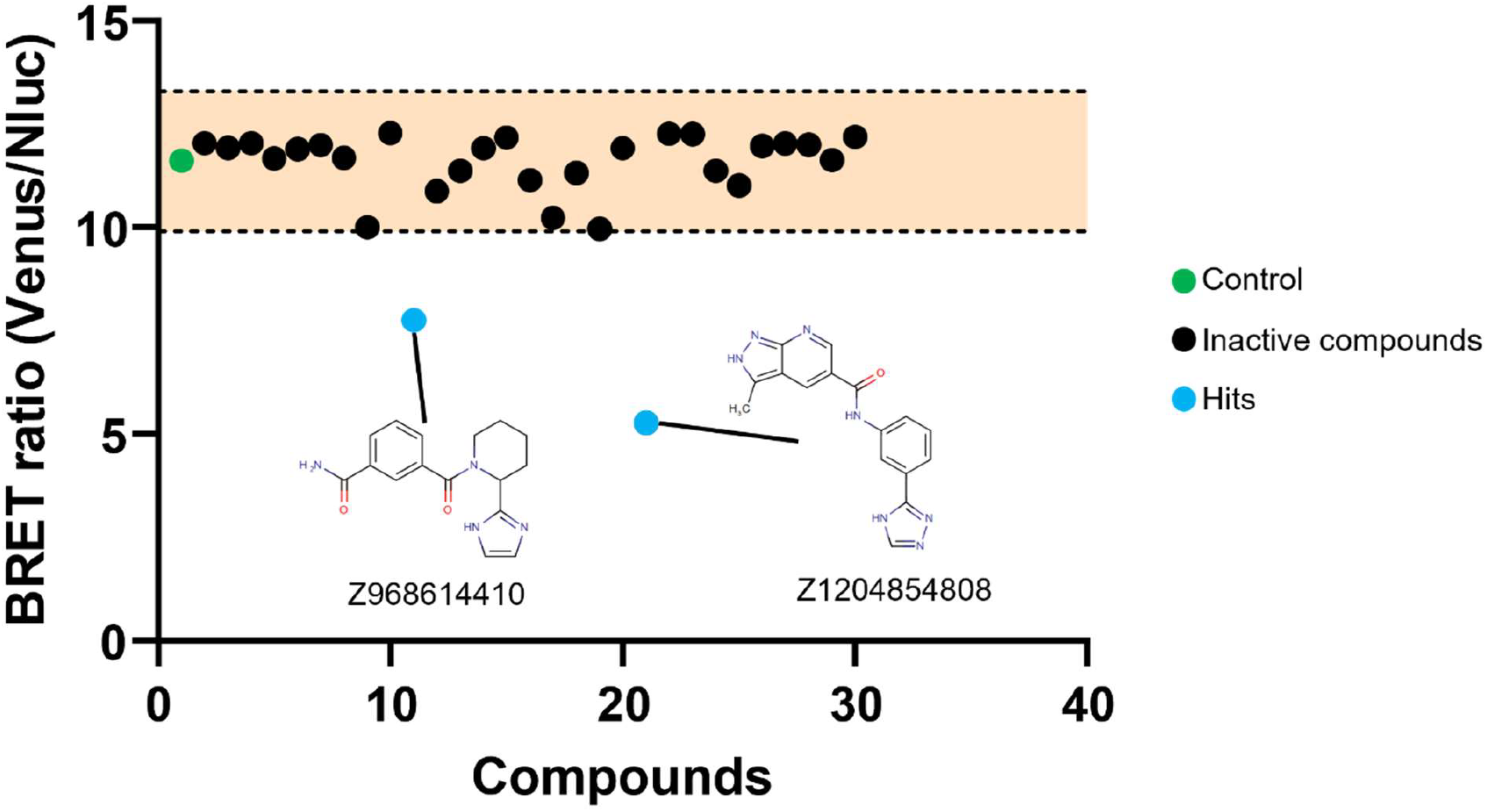
Functional screening of ensemble-predicted compounds using a live-cell NanoBRET assay. Twenty-nine compounds identified from ensemble-based virtual screening were evaluated for their ability to disrupt the NOS1-NOS1AP interaction in live cells. Compounds exhibiting reduced NanoBRET signal indicate functional disruption of the protein-protein interaction. Data are presented as mean ± SEM (n = 3).

To assess the ability of these compounds to inhibit the NOS1-NOS1AP interaction in a cellular context, we developed a NanoBRET-based assay system. In this system, NanoLuciferase was fused to the N-terminus of NOS1, while Venus fluorescent protein was tagged to the C-terminus of NOS1AP. When NOS1 and NOS1AP interact, energy transfer occurs from NanoLuciferase to Venus, producing a measurable BRET signal. Disruption of this interaction by a compound would result in a decrease in the NanoBRET ratio.

Among the three compounds tested, Z1204854808 (from here will be referred to as MC-3) demonstrated the most pronounced inhibitory effect on the NOS1-NOS1AP interaction (Figure 4). At the highest concentration tested, **MC-3** reduced the NanoBRET signal to approximately 57.5 ± 1.2% of the control, indicating substantial disruption of the protein-protein interaction. In comparison, Z968614410 and Z2607801368 showed relatively modest effects, reducing the signal to approximately 72.2 ± 0.6 % and 64.4 ± 3.7 % of control, respectively. The dose-dependent reduction in NanoBRET signal observed with **MC-3** suggested a specific and concentration-dependent inhibition of the NOS1-NOS1AP interaction.

**Figure 4.**
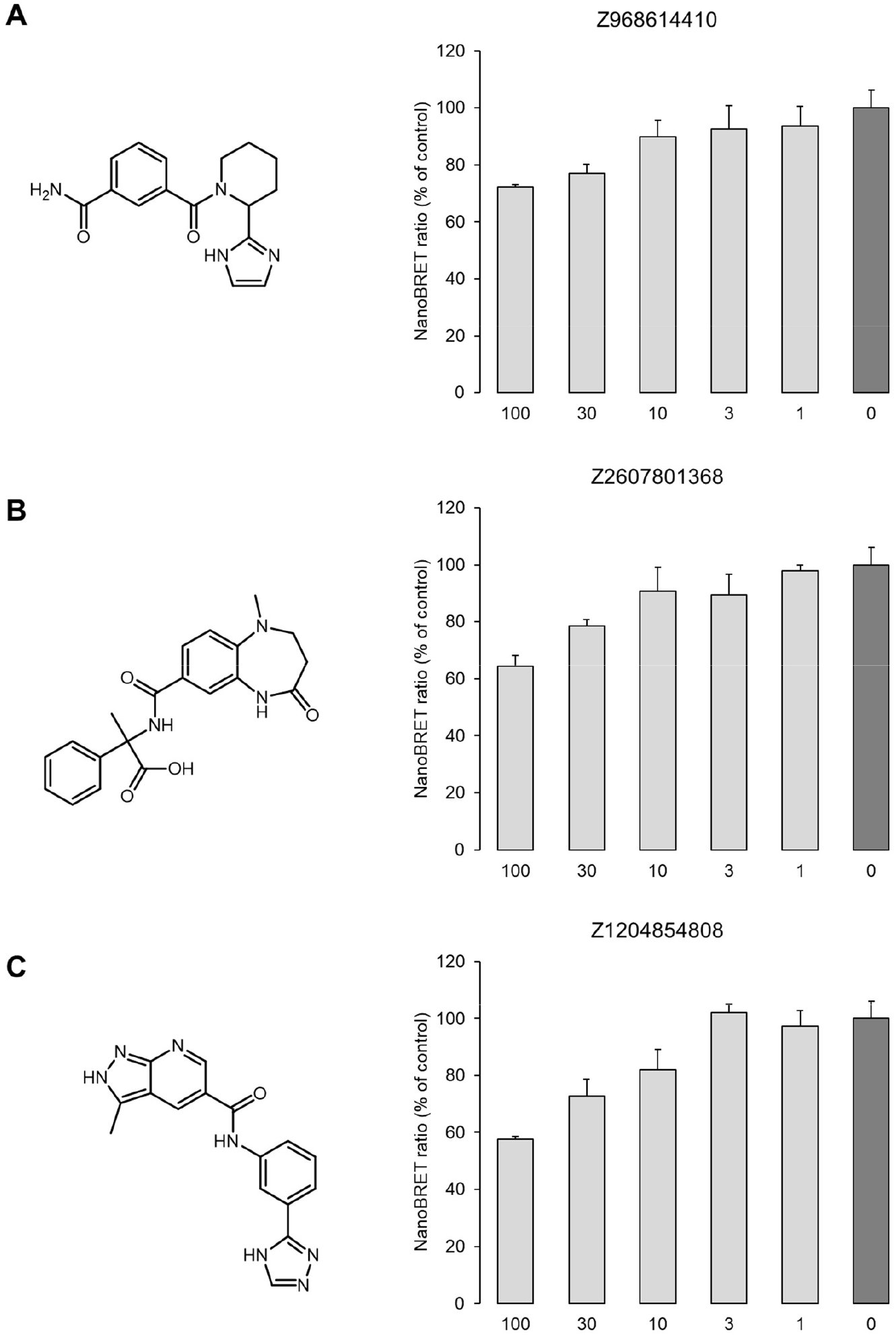
Identification of MC-3 as a NOS1-NOS1AP interaction inhibitor using NanoBRET assay. (A-C) Three hit compounds identified from in silico screening. Left panels show the chemical structures of each compound, and right panels show the concentration-dependent effects on NOS1-NOS1AP interaction assessed by NanoBRET assay. NanoBRET assay was performed using CHO-K1 cells co-expressing NanoLuciferase-tagged NOS1 (N-terminal) and Venus-tagged NOS1AP (C-terminal). Cells were treated with indicated concentrations (0, 1, 3, 10, 30, and 100 μM) of each compound, and the NanoBRET ratio was measured and expressed as a percentage of the vehicle control. (A) Z968614410, (B) Z2607801368, (C) **MC-3**. Among the tested compounds, **MC-3** exhibited the most potent inhibitory effect on the NOS1-NOS1AP interaction. Data are presented as mean ± SEM (n = 3).

The identification of **MC-3** highlights the importance of explicitly accounting for conformational heterogeneity when targeting dynamic protein–protein interaction interfaces. Because the nNOS PDZ domain populates at least two structurally distinct conformational states in solution, ligands optimized against a single static structure are unlikely to engage all functionally relevant binding geometries. The ability of **MC-3** to disrupt the NOS1-NOS1AP interaction in live cells, together with its direct engagement of the nNOS PDZ domain, indicates that it exploits structural features present in ligand-competent conformational states captured by the ensemble-based screening approach. Notably, inclusion of a minor but structurally distinct conformational state during virtual screening likely expanded the accessible chemical space and enabled identification of interaction modes that would be inaccessible in single-structure docking workflows. Collectively, these findings demonstrate how ensemble-aware screening can reveal small molecule modulators of protein-protein interactions that are otherwise refractory to conventional structure-based discovery strategies.

### 2.5. Confirmation of direct binding to NOS1 PDZ domain

To verify that **MC-3** directly binds to NOS1 and to identify its binding site, we performed a biophysical binding assay using spectral shift assay (Figure S2). This method enables the detection of ligand-protein interactions by monitoring changes in the intrinsic fluorescence properties of proteins upon compound binding. For this analysis, we utilized a recombinant protein containing only the PDZ domain of NOS1, which is known to mediate the interaction with NOS1AP.

The spectral shift analysis revealed a clear concentration-dependent change in the fluorescence ratio (670 nm/650 nm) at higher concentrations of **MC-3**, with a pronounced spectral shift observed at concentrations above 100 μM, indicating that the compound binds to the NOS1 PDZ domain at these concentrations (Figure S2). This finding is consistent with our NanoBRET results, where **MC-3** showed effective inhibition of the NOS1-NOS1AP interaction at similar concentration ranges, further supporting the direct binding of **MC-3** to the NOS1 PDZ domain as its mechanism of action. Taken together, these findings establish **MC-3** as a promising lead compound for further development as a NOS1-NOS1AP interaction inhibitor.

### 2.6. Selectivity assessment via CAPON binding assays

To test whether any of these small molecules interact directly with CAPON (NOS1AP), we performed a binding screen using the Dianthus Temperature-Related Intensity Change (TRIC) assay. Figure 5 summarizes the normalized fluorescence (Fnorm) values obtained across all compounds relative to the reference controls (protein plus dye only). Each compound was tested at a concentration of 200 nM against 10 nM his-tagged CAPON protein. Most compounds clustered within the established reference range (shaded area), which represents three times the standard deviation of the reference average, indicating no detectable target engagement under the assay conditionsThree compounds (Figure 5) showed clear deviations from this range, suggesting possible changes in the thermal stability of CAPON consistent with potential binding. Notably, two compounds produced marked reductions in Fnorm, pointing to possible ligand–protein interactions. These results served as an initial filter to identify candidates for follow-up biophysical validationTo further evaluate these preliminary hits, we assessed the two candidate compounds (Figure S3) using microscale thermophoresis (MST). Fluorescently labeled His-tagged CAPON protein was titrated with serial dilutions of both candidate compounds under optimized buffer (PBST) conditions containing 2% DMSO. The resulting thermophoretic responses were analyzed using the standard Hill model to calculate dissociation constants (Kd). Neither compound produced a clear dose-dependent change in normalized fluorescence. The scatter of the data points indicated that the thermophoretic response was insufficient to generate a reliable binding curve. Consequently, no dissociation constant (Kd) could be determined for either compound. These findings suggest that the apparent signals observed in the initial Dianthus screen do not translate into detectable binding by MST, indicating that none of the tested compounds is a CAPON binder under the conditions examined.

**Figure 5.**
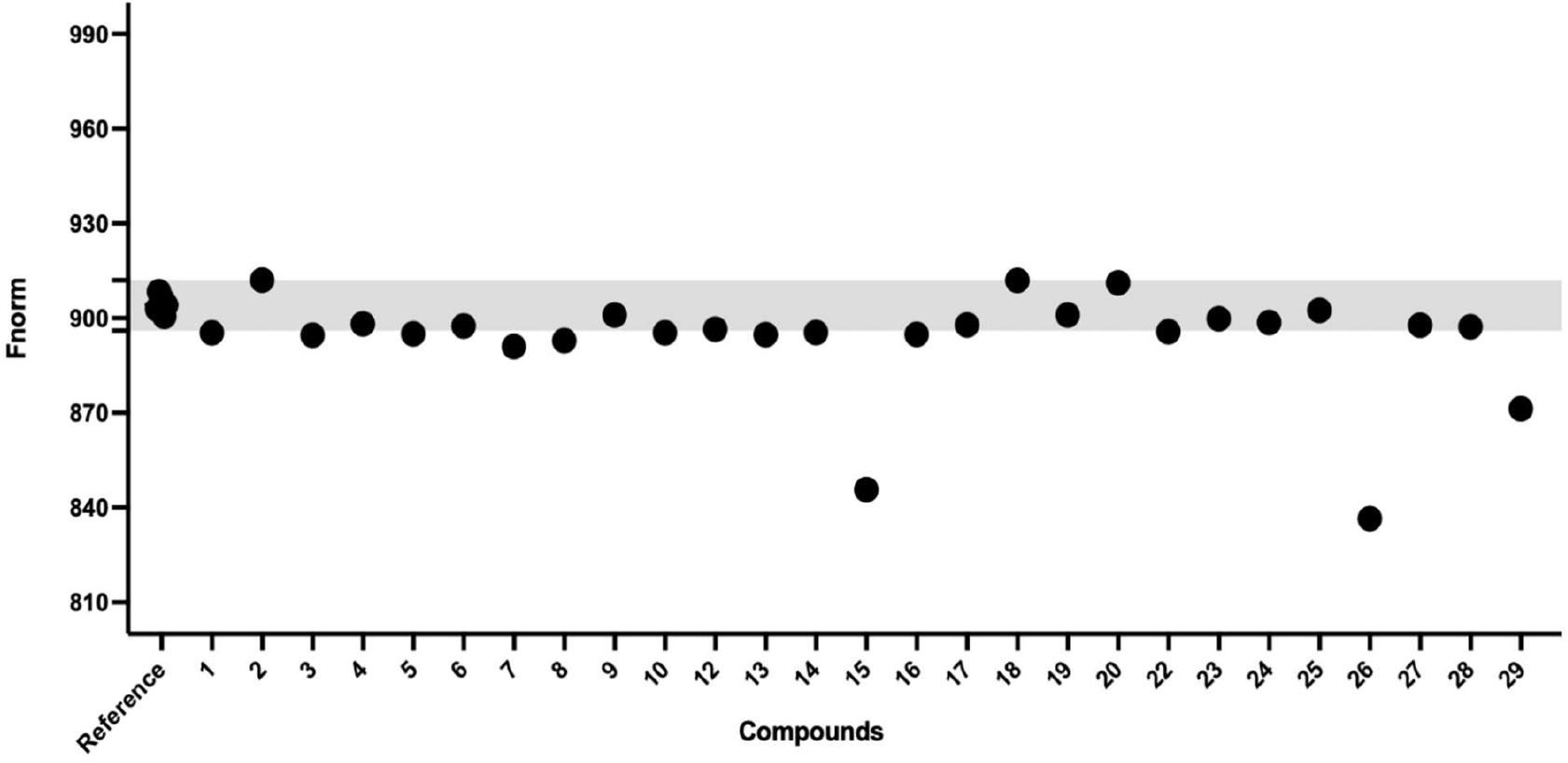
Assessment of direct compound binding to CAPON via MST. Normalized fluorescence (Fnorm) values are shown for all tested compounds relative to the reference control (CAPON protein plus dye only).

### 2.7. PK. Profiling

**MC-3** was profiled using a panel of in vitro physicochemical and ADME assays to evaluate its developability as a CNS-targeted small molecule candidate. Our physicochemical property assessment (Table 1) indicated moderate lipophilicity (cLogD_7.4_ = 1.2), consistent with the heteroaromatic nature of the scaffold and the presence of multiple hydrogen-bond donors and acceptors. Notably, **MC-3** exhibited aqueous kinetic solubility (78.3 µM at pH7.4), supporting its use in cell-based assays and enabling formulation at concentrations relevant for pharmacological evaluation.

**Table 1.**
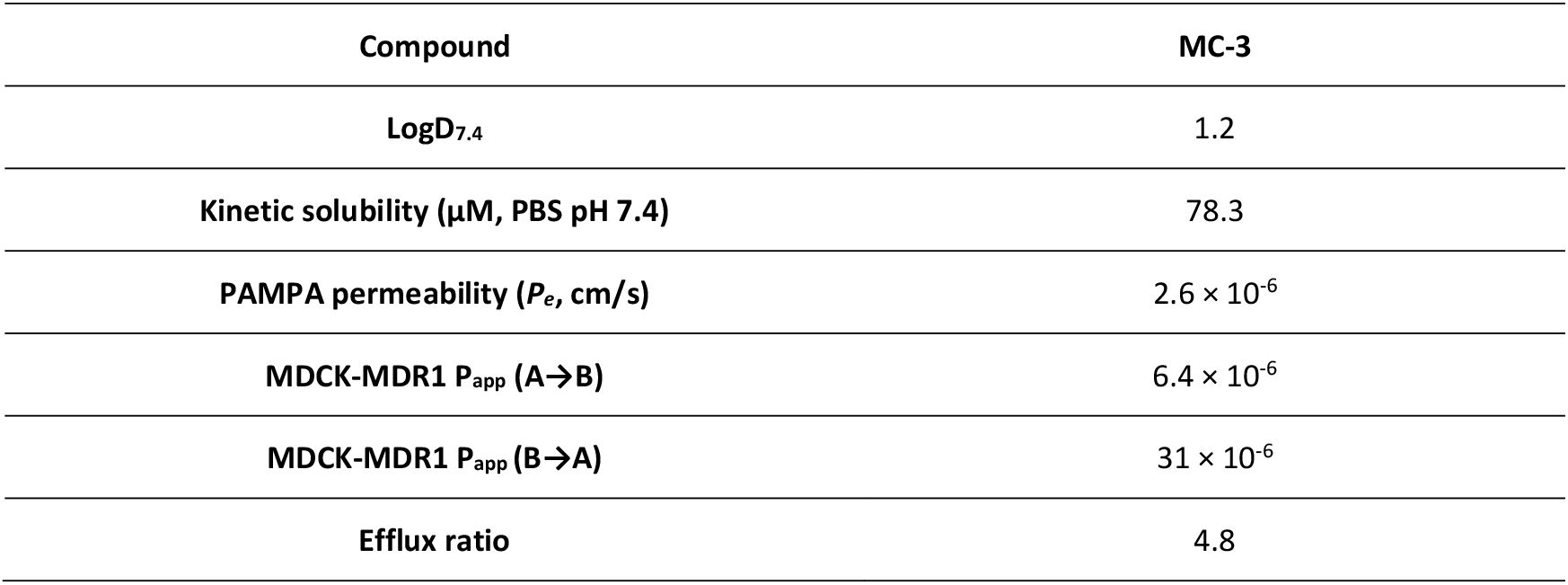
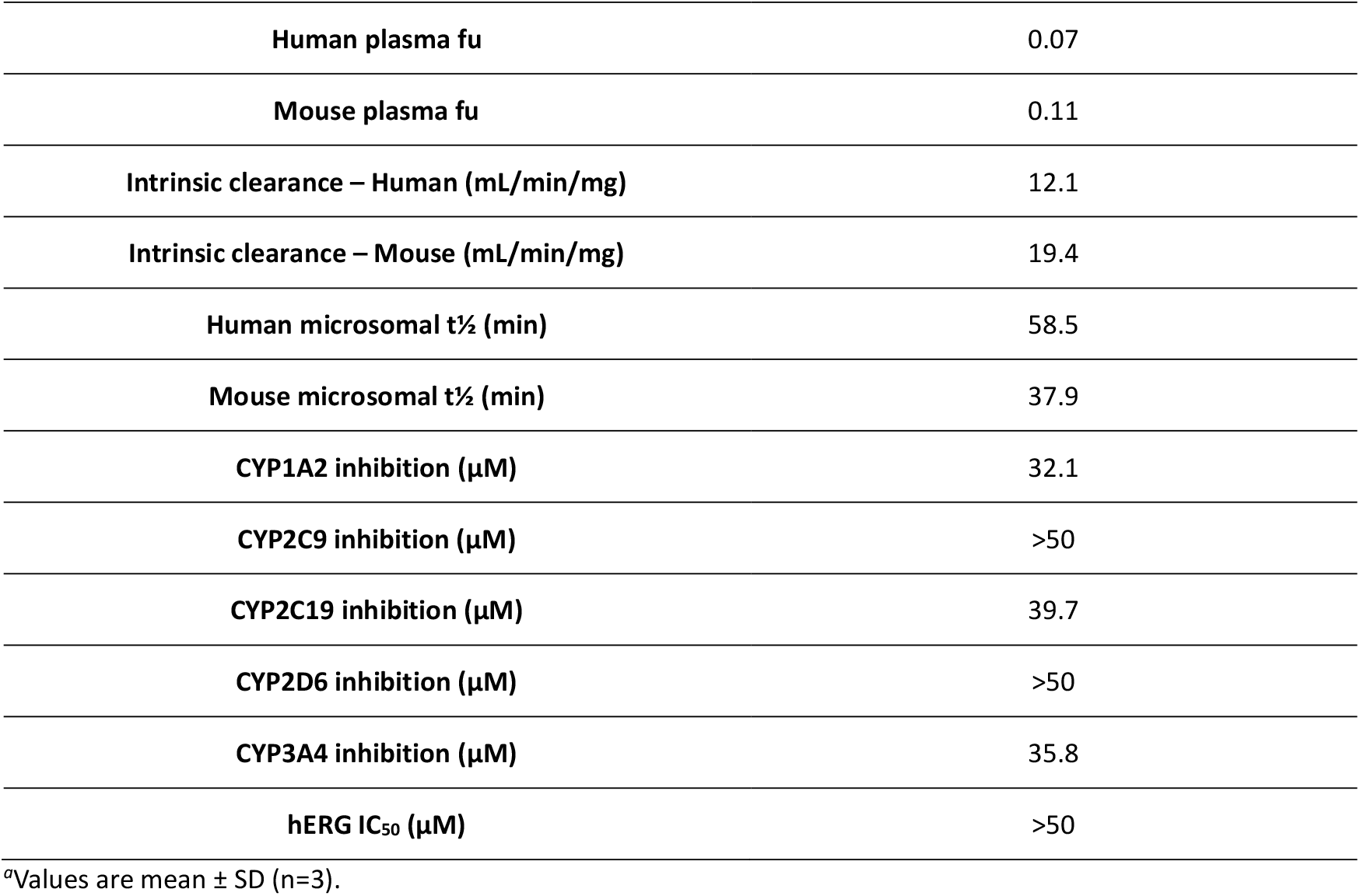
Assessment of the in vitro PK properties of **MC-3**.^a^.

| Compound | MC-3 |
| --- | --- |
| LogD<sub>7.4</sub> | 1.2 |
| Kinetic solubility (μM, PBS pH 7.4) | 78.3 |
| PAMPA permeability ($P_e$, cm/s) | $2.6 \times 10^{-6}$ |
| MDCK-MDR1 $P_{app}$ (A→B) | $6.4 \times 10^{-6}$ |
| MDCK-MDR1 $P_{app}$ (B→A) | $31 \times 10^{-6}$ |
| Efflux ratio | 4.8 |
| Human plasma fu | 0.07 |
| Mouse plasma fu | 0.11 |
| Intrinsic clearance – Human (mL/min/mg) | 12.1 |
| Intrinsic clearance – Mouse (mL/min/mg) | 19.4 |
| Human microsomal t<sub>1/2</sub> (min) | 58.5 |
| Mouse microsomal t<sub>1/2</sub> (min) | 37.9 |
| CYP1A2 inhibition (μM) | 32.1 |
| CYP2C9 inhibition (μM) | >50 |
| CYP2C19 inhibition (μM) | 39.7 |
| CYP2D6 inhibition (μM) | >50 |
| CYP3A4 inhibition (μM) | 35.8 |
| hERG IC<sub>50</sub> (μM) | >50 |

<sup>a</sup>Values are mean ± SD (n=3).

In vitro permeability assessments (Table 1) revealed acceptable passive diffusion across an artificial BBB membrane (PAMPA-BBB P_e_ = 2.6 × 10^−6^ cm/s). Consistent with its physicochemical profile, **MC-3** displayed moderate apparent permeability in MDCK-MDR1 cells (A→B P_app_ = 6.4 × 10^−6^ cm/s) accompanied by a higher B→A flux (31 × 10^−6^ cm/s), yielding an efflux ratio of 4.8. These data suggest that active efflux may contribute to limiting cellular accumulation and may influence CNS exposure, highlighting efflux liability as a key consideration for future optimization efforts. Protein binding studies indicated moderate to high plasma binding, with unbound fractions of 7% and 10% in human and mouse plasma, respectively (Table 1). Together, these data suggest that free drug exposure, rather than total concentration, will be the most relevant driver of biological activity.

Metabolic stability was assessed in human and mouse liver microsomes, where **MC-3** demonstrated moderate intrinsic clearance (HLM Cl_int_ 12.1 µL/min/mg; MLM Cl_int_ 19.4 µL/min/mg). These clearance values correspond to calculated in vitro microsomal half-lives of approximately 58.5 min in human and 37.9 min in mouse microsomes, indicating a scaffold with sufficient metabolic stability to support further PK evaluation while retaining room for optimization if needed. Importantly, **MC-3** showed minimal inhibition of major cytochrome P450 isoforms (IC_50_ ≥ 30 µM across the panel) and no detectable hERG liability (IC_50_ > 50 µM), supporting a favorable early safety and drug-drug interaction profile.

Overall, the in vitro ADME profile of **MC-3** reflects a balanced profile, characterized by moderate solubility, acceptable metabolic stability, and permeability that is likely influenced by both polarity and transporter-mediated efflux. These features are consistent with an early lead candidate and provide the basis for structure-guided optimization, including reduction of efflux susceptibility and improving solubility, while preserving target engagement and cellular activity relevant to AD.

### 2.8. Biological validation of MC-3 as a drug candidate for AD

To evaluate the therapeutic potential of small molecule CAPON inhibition in AD, we examined the effects of the lead compound **MC-3** across three complementary cellular assays probing amyloid-driven neurotoxicity, CAPON-dependent nitrosative signaling, and tau pathology. Together, these assays interrogate convergent pathological pathways central to neurodegeneration.

Exposure of primary cortical neurons to Aβ_42_ oligomers resulted in a robust increase in neuronal cytotoxicity, as quantified by LDH release following 24 h incubation (Figure 6A). Pretreatment with **MC-3** produced a clear, concentration-dependent neuroprotective effect at 10, 25, and 50 µM, progressively reducing Aβ-induced toxicity and restoring neuronal viability toward basal levels (Figure 6A). At the highest concentration tested, **MC-3** nearly normalized LDH release to that observed in untreated controls. Importantly, **MC-3** did not induce detectable cytotoxicity when administered alone, indicating that the observed protection was not attributable to nonspecific effects on neuronal survival. These results demonstrate that CAPON inhibition by **MC-3** effectively protects neurons from amyloid-mediated injury, a central pathogenic event in AD.

**Figure 6.**
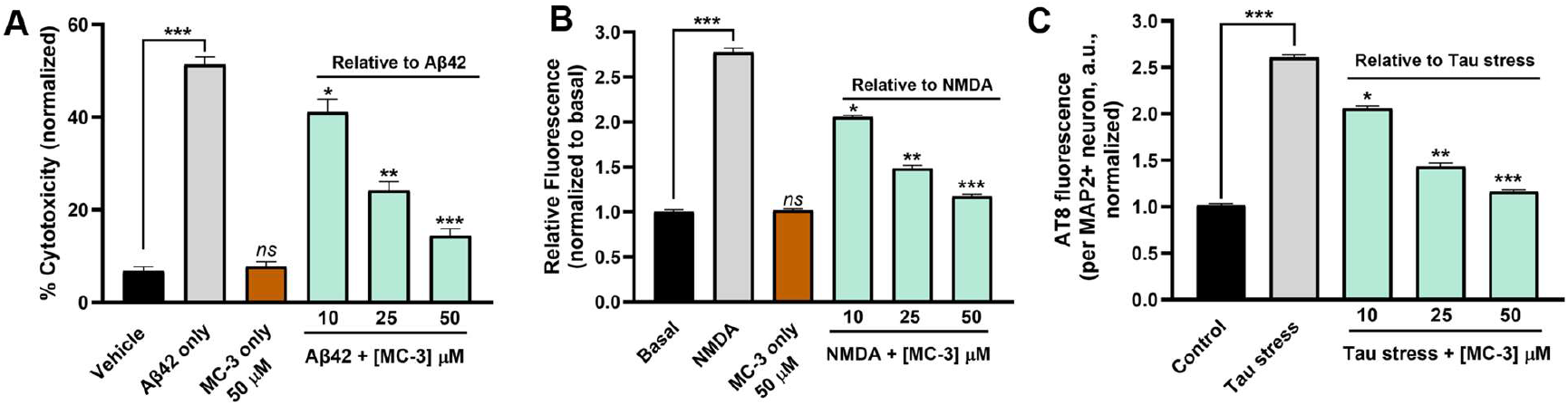
MC-3 engages CAPON-dependent pathways to protect neurons from AD-relevant pathways. **(A) MC-3** protects primary cortical neurons from Aβ42-induced cytotoxicity. Pretreatment with **MC-3** (10, 25, or 50 µM) produced a concentration-dependent reduction in LDH release following Aβ42 oligomer exposure. **(B) MC-3** suppresses pathological NMDA-driven nitric oxide signaling. **MC-3** pretreatment attenuated NMDA-induced intracellular NO production, measured by DAF-FM DA fluorescence, in a dose-dependent manner. **(C) MC-3** reduces pathological tau phosphorylation in neuronal cells. Neurons subjected to tau-stress conditions were treated with **MC-3** (10, 25, or 50 µM), and tau pathology was assessed by AT8 immunofluorescence. Quantitative analysis revealed a concentration-dependent decrease in AT8 intensity per MAP2^+^ neuron, normalized to control. Data represent mean ± SEM from three independent measurements. * p < 0.05, ** p < 0.01, *** p < 0.001, and (ns) denotes nonsignificant relative to untreated control.

To determine whether the neuroprotective effects of **MC-3** were associated with modulation of CAPON-dependent signaling downstream of NMDA receptor activation, we measured neuronal nitric oxide production using the fluorescent probe DAF-FM DA (4-amino-5-methylamino-2’,7’-difluorofluorescein diacetate). Brief NMDA stimulation elicited a pronounced increase in intracellular NO levels, consistent with activation of nNOS-dependent nitrosative signaling. **MC-3** treatment significantly attenuated NMDA-induced NO production in a dose-dependent manner, with partial suppression observed at 10 µM and near-basal NO levels achieved at higher concentrations (Figure 6B). **MC-3** alone did not affect basal NO production, suggesting that CAPON inhibition selectively dampens pathological NMDA-driven signaling rather than globally suppressing neuronal activity. Because nitric oxide generation represents a proximal functional output of nNOS activation, these findings indicate that **MC-3** effectively disrupts CAPON-mediated coupling between NMDA receptors and nNOS. This suppression of nitrosative stress provides a mechanistic link between CAPON inhibition and protection against amyloid-induced neurotoxicity.

Given the reported role of CAPON in regulating tau phosphorylation, we next examined whether **MC-3** modulates pathological tau phosphorylation in neuronal cells. Under tau-stress conditions, neurons exhibited a marked increase in AT8 immunoreactivity, consistent with elevated phosphorylation of tau at disease-relevant epitopes. Treatment with **MC-3** resulted in a concentration-dependent reduction in AT8 fluorescence intensity per MAP2^+^ neuron (Figure 6C). Notably, **MC-3** reduced AT8 signal without affecting neuronal counts or MAP2 labeling, indicating that the observed effects were not attributable to nonspecific neuronal loss. These results demonstrate that **MC-3** attenuates pathological tau phosphorylation in neuronal cells.

Collectively, these findings demonstrate that pharmacological inhibition of CAPON using **MC-3** exerts convergent protective effects across the amyloid, nitrosative stress, and tau axes of AD pathology. By suppressing NMDA-dependent nitric oxide signaling, **MC-3** dampens a key neurotoxic pathway that links excitotoxicity to both amyloid-mediated neuronal injury and tau pathology. These data position CAPON as a compelling intracellular target for small molecule intervention in AD and establish **MC-3** as a promising lead compound capable of addressing multiple pathological drivers of neurodegeneration.

## 3. Methods

### 3.1. NMR conformations analysis

The 1B8Q multi-model NMR ensemble was analyzed using AtlasNMR to quantify backbone structural heterogeneity and identify representative conformations for ensemble-based virtual screening. All models were superimposed onto a reference frame (Model 1) using least-squares fitting of backbone atoms (Cα, C, N, O) within chain A to minimize translational and rotational variance. Following structural alignment, pairwise root mean square deviation (RMSD) matrices were computed across all 508 backbone atoms for each model pair, generating a comprehensive 15×15 distance matrix that captured the complete conformational landscape of the ensemble. Additional structural metrics were calculated, including per-model radius of gyration (Rg), per-residue root mean square fluctuation (RMSF), and global mean RMSD, to quantify both individual model compactness and ensemble-wide flexibility patterns.

Conformational outliers were identified using a consensus approach integrating three independent methods: (i) interquartile range (IQR) analysis on mean RMSD distributions with thresholds defined as Q1 - 1.5×IQR to Q3 + 1.5×IQR, (ii) Isolation Forest algorithm applied to flattened coordinate matrices with a contamination parameter of 0.1, and (iii) hierarchical clustering with Ward’s linkage to detect singleton clusters at μ + 2σ distance thresholds. Models flagged by at least two of the three methods were designated consensus outliers and excluded from subsequent representative structure selection.

The optimal number of clusters (k) was determined by systematically evaluating k = 2–6 using multiple quality metrics on non-outlier structures: silhouette coefficient (maximized), Davies-Bouldin index (minimized), Calinski-Harabasz score (maximized), and elbow method analysis of within-cluster sum of squares (inertia). Final clustering was performed using k-means with k = 2, initialized with 20 independent runs to ensure convergence to the global optimum. Cluster quality was validated through silhouette analysis, while intra-cluster diversity was quantified using mean pairwise RMSD, cluster diameter (maximum pairwise distance), and compactness scores.

Representative structures for each cluster were identified using a consensus voting scheme that integrated four independent selection criteria: (i) medoid selection based on minimum average RMSD to all cluster members, (ii) centroid proximity calculated as minimum Euclidean distance to geometric mean coordinates, (iii) minimax optimization identifying the structure with minimum maximum pairwise RMSD, and (iv) density peak detection based on maximum neighbor count within the 25th percentile RMSD threshold. The model most frequently selected across all four methods was designated as the primary cluster representative. Representative quality was assessed using a composite scoring function with the following weightings: 35% normalized average RMSD (lower is better), 25% normalized maximum RMSD (lower is better), 25% coverage (percentage of cluster members within 2 Å RMSD), and 15% centrality (inverse distance rank within cluster). Inter-representative RMSD was calculated to assess conformational space coverage between selected representatives, and principal component analysis (PCA) was performed on flattened coordinate matrices to visualize cluster separation in reduced-dimensionality space. All computational analyses were implemented in Python 3.9 using scikit-learn v1.0 for clustering and dimensionality reduction, SciPy v1.7 for hierarchical clustering methods, and custom algorithms for composite quality scoring and consensus ranking.

### 3.2. Structure-Based Virtual Screening and Consensus Scoring

Structure-based virtual screening was applied to identify potential CAPON-NOS interaction modulators from the Enamine compound libraries using a hierarchical docking approach. The NOS1 protein structure was obtained from the Protein Data Bank (PDB ID: 1B8Q^23^) and prepared for virtual screening using the Protein Preparation Wizard in Schrödinger Suite. The CAPON-NOS binding site was identified and validated through analysis of the co-crystal structure, with receptor grids generated using Glide’s grid generation module to represent the shape and electrostatic features of the binding pocket through multiple separate field sets.

The consensus virtual screening protocol employed a three-stage hierarchical filtering approach using Glide’s High-Throughput Virtual Screening (HTVS), Standard Precision (SP), and Extra Precision (XP) docking modes applied to two representative protein conformations (models 3 and 13) to account for receptor flexibility according to our previously published protocol^24^.

### 3.3. NanoBRET assay for NOS1-NOS1AP interaction

To monitor the NOS1-NOS1AP protein-protein interaction in living cells, a NanoBRET-based assay system was established. The full-length human NOS1 was cloned with NanoLuciferase (NanoLuc) fused to its N-terminus, while full-length human NOS1AP (CAPON) was cloned with Venus fluorescent protein fused to its C-terminus. CHO-K1 cells were co-transfected with both constructs using an appropriate transfection reagent. Transfected cells were then seeded in white 96-well plates and cultured until reaching approximately 80% confluency. Prior to compound treatment, growth media was removed, and cells were washed twice with PBS, followed by the addition of HBSS solution. Cells were then treated with the indicated concentrations of test compounds or vehicle control (DMSO) for 3 hours. NanoBRET measurements were performed by adding NanoLuc substrate and measuring donor emission (460 nm) and acceptor emission (535 nm) using a Tecan plate reader. The NanoBRET ratio was calculated as the acceptor/donor emission ratio and normalized to the vehicle control. All experiments were performed in triplicate and repeated at least three times independently.

### 3.4. Spectral shift binding assay

To confirm the direct binding of compounds to the NOS1 PDZ domain or NOS1AP, spectral shift measurements were performed using MonolithX (NanoTemper Technologies, Munich, Germany). Recombinant NOS1 PDZ domain protein (Abbexa, Cambridge, UK) was labeled with a fluorescent dye using the Spectral Shift Optimized Protein Labeling Kit - For His-Tag according to the manufacturer’s protocol. PBS containing 0.005% Tween was used as the assay buffer. For the binding assay, a constant concentration of labeled protein was mixed with the indicated concentrations of **MC-3** and incubated at room temperature for 30 minutes. Samples were then loaded into capillaries, and spectral shift was measured by monitoring the change in the fluorescence emission ratio (670 nm/650 nm) using the MonolithX instrument. The binding curve was generated by plotting the fluorescence ratio against the logarithm of compound concentration.

### 3.5. Single-dose Dianthus screening

Protein-ligand binding was initially evaluated by microscale thermophoresis (MST). Recombinant CAPON carrying a C-terminal His tag was fluorescently labeled using the RED-tris-NTA dye (NanoTemper Technologies) following the manufacturer’s protocol. 200 nM protein was incubated with 100 nM dye in Phosphate Buffered Saline PBS supplemented with 0.05% Tween-20 buffer (pH 7.4) for 30 minutes at room temperature in the absence of light. Following labeling, the protein was diluted into PBST and combined with test compounds to yield final assay concentrations of 20 nM protein and 250 µM compound. Samples were incubated for 30 minutes at ambient temperature, briefly centrifuged, and subsequently analyzed on a Dianthus NT.23 Pico instrument. Assay buffer containing DMSO alone served as the negative control. Each measurement was performed in triplicate, and the reported values represent the averaged data.

### 3.6. Dose–response characterization using Microscale thermophoresis (MST)

Compounds that produced a clear binding response in the single-point screen were subsequently evaluated in a concentration-dependent MST assay. Twelve-point serial dilutions were prepared, spanning a starting concentration of 250 µM down to the low-nanomolar range, and mixed with labeled CAPON-His protein while maintaining a final DMSO concentration of 2%. After a 30-minute equilibration step at room temperature, samples were loaded into standard MST capillaries and measured on a Monolith NT.115 instrument, using medium-to-high infrared laser power and 60–80% LED excitation in the red detection channel. Dissociation constants (Kd) were calculated using the MO.Affinity Analysis package (NanoTemper Technologies).

### 3.7. Evaluation of PK and physicochemical properties

These experiments were conducted following our previously reported methods.^1^ The evaluations included LogD7.4 determination, microsomal stability, kinetic solubility, and cytotoxicity profiling across multiple cell lines. Solubility was assessed using UV-visible spectrophotometry.

### 3.8. Aβ_42_-Induced Neurotoxicity and Neuroprotection Assay

Primary cortical neurons (from E18 rat embryos) were cultured on poly-D-lysine–coated plates in Neurobasal medium supplemented with B27 and GlutaMAX. Neurons were used between DIV7-9. For Aβ_42_ oligomer preparation, Aβ_42_ peptide was dissolved in HFIP, dried to a film, resuspended in DMSO, diluted in phenol-red–free F12 medium, and incubated at 4 °C for 24 h. Neurons were pretreated with **MC-3** (10, 25, and 50 µM) or vehicle (0.1% DMSO) for 1 h prior to exposure to Aβ_42_ oligomers. Following 24 h incubation, neuronal cytotoxicity was quantified using an LDH release assay. Maximum LDH release was determined using lysis buffer-treated wells, and background was subtracted using no-cell controls. In selected experiments, neuronal viability was confirmed using complementary live/dead staining or ATP-based viability assays.

### 3.9. NMDA-Induced Nitrosative Stress Signaling

Primary cortical neurons (DIV8) were pretreated with **MC-3** (10, 25, and 50 µM) or vehicle for 1 h, followed by brief NMDA stimulation (50 µM, 10 min) to activate nNOS-dependent signaling. Cells were then washed and processed immediately. Nitric oxide (NO) production was quantified using the fluorescent probe DAF-FM DA, loaded into neurons prior to NMDA stimulation. Fluorescence intensity was measured by a Tecan plate reader.

### 3.10. Modulation of Tau Pathology in Neuronal Cells

Primary cortical neurons were cultured as described above and used between DIV7-10. To induce tau pathology (tau-stress), neurons were treated with okadaic acid (OA, 25 nM) for 24 h, to promote tau hyperphosphorylation through inhibition of protein phosphatases. Neurons were treated with **MC-3** (10, 25, or 50 µM) or vehicle control for 24 h.

Following treatment, cells were fixed with 4% paraformaldehyde, permeabilized, and blocked using standard conditions. Neurons were incubated with an antibody against phosphorylated tau (AT8; Ser202/Thr205) together with an antibody against MAP2 to label neuronal cell bodies and neurites. Nuclei were counterstained with DAPI. Fluorescent secondary antibodies were applied, and cells were imaged using a Tecan Spark. MAP2-positive neurons were identified, and AT8 fluorescence intensity was quantified on a per-neuron basis. AT8 signal was normalized to neuronal area and expressed as mean AT8 intensity per MAP2^+^ neuron. Data were normalized to control conditions and are reported as mean ± SEM from three independent experiments.

## 4. Conclusions

In summary, this work demonstrates that explicitly incorporating experimentally observed conformational heterogeneity can enable small molecule discovery against dynamic protein-protein interaction interfaces that are poorly served by single-structure approaches. By converting NMR ensemble data into screening-ready conformational hypotheses, the AtlasNMR framework provides a generalizable strategy for exploiting protein dynamics in chemical discovery. Application of this approach to the CAPON/nNOS system enabled identification of **MC-3**, a small molecule modulator that disrupts the NOS1-NOS1AP interaction and confers convergent neuroprotective effects across multiple pathological axes relevant to Alzheimer’s disease. More broadly, this ensemble-aware strategy offers a practical path for targeting conformationally plastic interaction domains and establishes a general chemical framework for exploiting experimentally observed protein dynamics in small molecule discovery.

## ASSOCIATED CONTENT

### Supporting Information

The Supporting Information is available free of charge on the ACS Publications website.

Conformational clustering and structural analysis of the nNOS PDZ domain NMR ensemble and Direct binding of **MC-3** to the NOS1 PDZ domain confirmed by spectral shift assay (PDF)

## AUTHOR INFORMATION

### Author Contributions

The manuscript was written through contributions of all authors. All authors have given approval to the final version of the manuscript.

## Acknowledgment

We acknowledge funding from the National Institutes on Aging (NIA) RF1AG084635 (PI: Gabr).

---

Insert Table of Contents artwork here

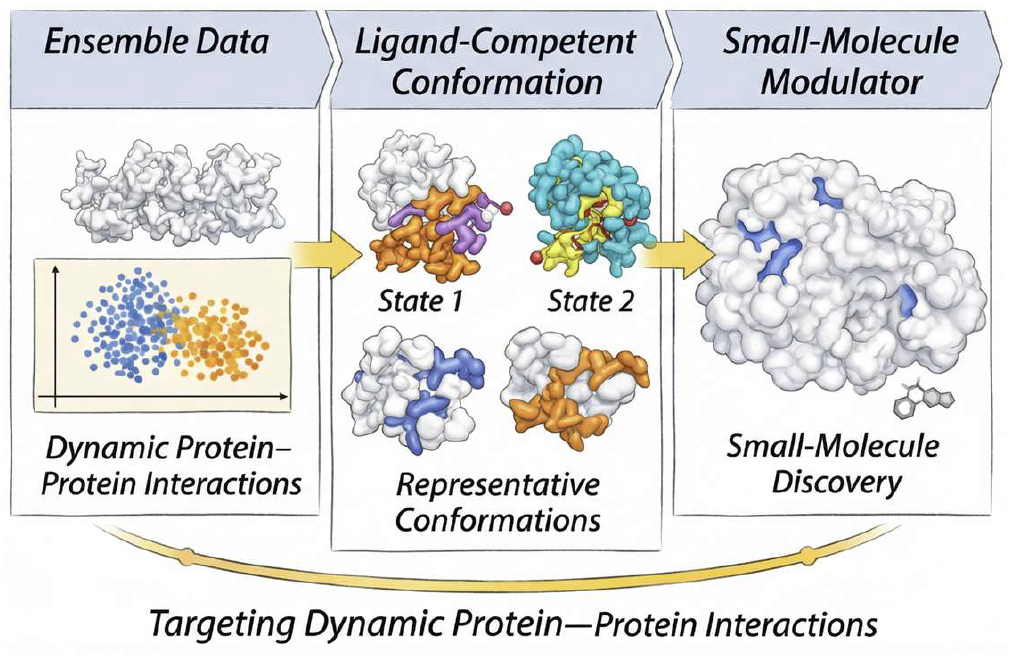

