## Supporting Information for "Exploiting NMR Ensemble Heterogeneity Enables Small Molecule Discovery Against Dynamic Protein-Protein Interfaces"

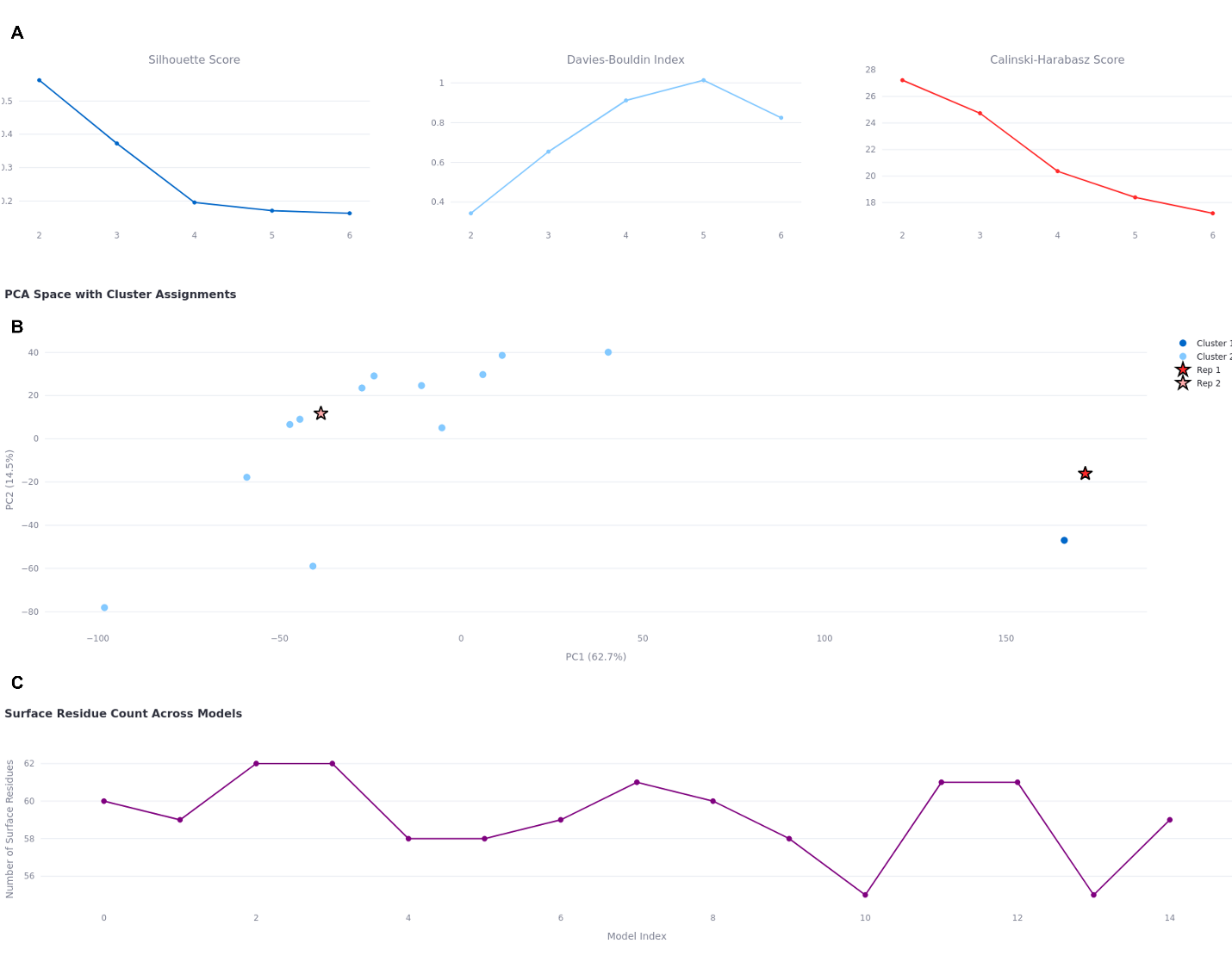

**Figure S1. Conformational clustering and structural analysis of the nNOS PDZ domain NMR ensemble.**
(A) Cluster validation metrics used to determine the optimal number of conformational states in the NOS/CAPON NMR ensemble (PDB ID: 1B8Q), analyzed using AtlasNMR. The silhouette coefficient, Davies–Bouldin Index, and Calinski–Harabasz score were evaluated across different values of *k*. (B) Principal component analysis (PCA) of the NMR ensemble with cluster assignments projected onto the first two principal components. The two clusters (C0 and C1) are clearly separated along PC1, reflecting a dominant conformational transition. Representative conformers 3 (C0) and 13 (C1), highlighted by star symbols, occupy opposite extremes of PC1 and capture the maximal structural diversity of the ensemble. (C) Surface-accessible residue counts across the NMR ensemble.

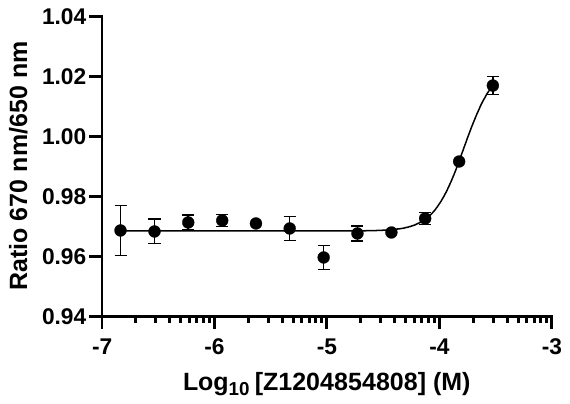

**Figure S2. Direct binding of MC-3 to the NOS1 PDZ domain confirmed by spectral shift assay.** Binding of **MC-3** to the recombinant NOS1 PDZ domain protein was assessed using spectral shift assay. The change in the fluorescence emission ratio (670 nm/650 nm) was measured across a range of compound concentrations. The concentration-dependent spectral shift observed at higher concentrations indicates direct binding of **MC-3** to the NOS1 PDZ domain. Data are presented as mean ± SEM (n = 3).

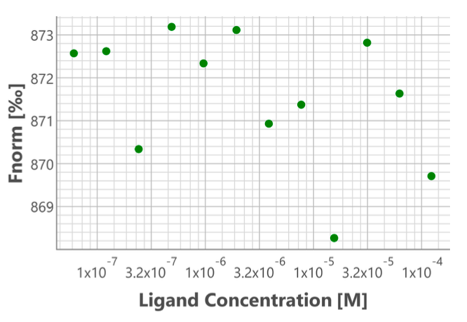

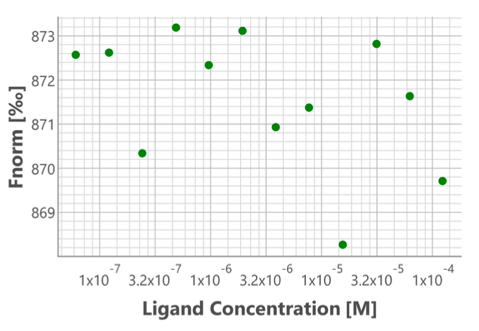

**Figure S3. MST analysis of the two hits predicted to bind to CAPON from the selectivity assay.**
